# A comprehensive AMR genotype-phenotype database (CABBAGE)

**DOI:** 10.1101/2025.11.12.688105

**Authors:** Emily Dickens, Romain Derelle, Robert Beardmore, Anita Suresh, Swapna Uplekar, Andrey G Azov, Tatiana A Gurbich, Bilal El Houdaigui, Jon Keatley, Sofiia Ochkalova, Orges Koci, Nadim M Rahman, Anu Shivalikanjli, Andrea Winterbottom, Galabina Yordanova, Helen Parkinson, Andrew D Yates, Robert D Finn, John A Lees, Leonid Chindelevitch

## Abstract

Addressing the growing threat of antimicrobial resistance (AMR) requires the development of large-scale resources that link bacterial genomic data with phenotypic antimicrobial resistance profiles. Such datasets are essential for advancing genotype-based predictions of resistance to uncover novel resistance mechanisms, as well as identifying and tracking global trends. Here, we describe the development of the ‘Comprehensive Assessment of Bacterial-Based AMR prediction from GEnotypes’ (CABBAGE) database, linking bacterial genomes to associated antibiotic susceptibility data and relevant metadata across WHO Bacterial Priority Pathogens, sourced from both publications and existing databases, and curated into a format that is compatible with, and extends, both NCBI and ENA formats. The resulting CABBAGE database, comprising over 170,000 unique sequenced isolates and approximately 1.7 million genome-phenotype pairs linked to extensive metadata, represents the largest database of its kind, consolidating existing AMR phenotype-genotype data into a single unified format. CABBAGE encompasses a broad range of antimicrobials, facilitating the analysis of global resistance trends as well as benchmarks of genotype-to-phenotype predictive methods, and empowering further research uses. The database is freely accessible at https://www.ebi.ac.uk/amr and is currently being integrated with the BioSample database, enabling easy access for the AMR research community.

## 1 INTRODUCTION

Antimicrobial resistance poses a substantial risk to global health, resulting in increased morbidity, mortality, and healthcare associated costs. Addressing this challenge requires a multifaceted approach, ranging from clinical interventions such as enhanced antibiotic stewardship and diagnostics, to pharmaceutical initiatives, including the development of novel antimicrobials. Among these, advances in whole-genome sequencing (WGS) have raised the possibility of using genomics-based diagnostics alongside, or as a replacement for, traditional antibiotic susceptibility tests (ASTs). However, while numerous antimicrobial resistance genes have been identified, discordance between genotypic predictions and phenotypic resistance is observed across numerous studies [1, 2, 3]. A key step in addressing this discordance is the development of large-scale, curated datasets that integrate AMR phenotypes with corresponding genomic data, with the aim of characterising resistance mechanisms, including those not yet genetically defined.

The World Health Organisation (WHO) has identified a list of bacterial pathogens which are a priority for new antibiotic development, encompassing 26 bacterial pathogen-antibiotic combinations in 2017 [4], revised to 24 in 2024 [5]. However, while the development of novel antimicrobials remains critical in addressing the growing threat of AMR, this strategy is often viewed as unsustainable due to the substantial costs associated with drug development and the risk of resistance developing soon after new antimicrobials are introduced into clinical use. An alternative approach is to maximise the effectiveness of existing antimicrobials with appropriate diagnostics and antimicrobial stewardship efforts [6, 7]. This could involve making treatment decisions informed by aggregated phenotypic and genotypic AMR data, however, clinically-derived AST results are not routinely shared with the scientific community, and genomes are not sequenced as part of standard practice for the majority of pathogens. Nevertheless, a wealth of genotypic and phenotypic information relevant to AMR already exists in various formats within public repositories.

A number of AMR databases have been developed to support research and surveillance efforts. Key resources include the Bacterial and Viral Bioinformatics Resource Center (BV-BRC) [8], the NCBI BioSample Antibiograms database [9] and the National Database of Antibiotic Resistant Organisms (NDARO) [10], all of which provide extensive bacterial genomic data linked to antimicrobial susceptibility data. Similarly, PubMLST [11], although primarily focusing on the provision of MLST information, also integrates bacterial population genomic data with phenotypic information, while platforms such as Microreact [12] and PathogenWatch [13] enable the visualisation of global AMR data alongside accompanying genotypic data when available. National level surveillance is also supported by agencies in the U.S. such as the Centres for Disease Control and Prevention (CDC) [14], which reports AMR trends, and the National Antimicrobial Resistance Monitoring System (NARMS) [15], which focuses on foodborne bacteria. Additionally, the European COMPARE Consortium [16] has created an AMR data hub consisting of over 2,500 sequenced isolates linked to structured phenotypic data. However, despite the availability of these resources, current AMR data remains largely fragmented across various institutions and regions, limiting its collective influence. There is therefore a need to develop a single, harmonised AMR database that integrates existing data, standardises data formats, and incorporates expert curation. Publications often also contain genotype-phenotype data and metadata that has not been included in any of the aforementioned resources, and can therefore provide an additional valuable source of information on AMR phenotypes and matching genotypes. A key challenge is that they can rarely be processed in a fully automated way.

The work presented in this study aimed to combine all available isolates of species belonging to the WHO Priority Pathogens with both genomic and antibiotic susceptibility testing data, drawing on two sources: an analysis of existing databases containing such datasets, as well as additional large AMR genotype-phenotype datasets extracted from the literature, followed by a process of data standardisation and curation. The end result is the ‘Comprehensive Assessment of Bacterial-Based AMR prediction from GEnotypes’ (CABBAGE) database, comprising over 170,000 unique sequenced isolates and approximately 1.7 million genome-phenotype pairs linked to extensive metadata. Here, we present an overview of CABBAGE, including the availability of data among different antibiotics and pathogens, the spatio-temporal data distribution, as well as a comparative analysis with ATLAS [17], a global AMR dataset encompassing over 900,000 isolates with associated antimicrobial susceptibility data, but no genotypic data aside from the presence/absence of a limited number of *β*-lactamase genes. Lastly, we describe how to access the CABBAGE dataset via the Antimicrobial Resistance Portal at EMBL-EBI (https://www.ebi.ac.uk/amr).

CABBAGE is a valuable resource, with applications ranging from advancing genotype-to-phenotype machine learning models that further our understanding of AMR mechanisms, to improvements in clinical diagnostics and global AMR surveillance. Additionally, the Antimicrobial Resistance Portal encourages data sharing by data producers in a standardised format, facilitating the long-term growth of CABBAGE.

## 2 METHODS

### 2.1 Data collection

A literature review was carried out with the aim of identifying and extracting data from publications containing at least 100 sequenced isolates on the WHO Bacterial Priority Pathogens List, with antimicrobial resistance phenotypes available for at least one drug. Note that this 100 isolate threshold was adjusted for some pathogens to account for their over- or under-representation in the literature, as detailed in the Supplementary Methods (S1.1). For each dataset with publicly available data, tables containing phenotypes as well as the accession numbers for the genotypes were extracted and standardised into a CSV format. These files were then processed via custom scripts, available at https://github.com/Leonardini/CABBAGE, to identify all relevant information on the genotype, phenotype, and metadata.

In addition to the datasets extracted from the literature, we identified nine public databases (Microreact, NARMS, NDARO, Pathogenwatch, BV-BRC [formerly known as PATRIC], PubMLST, COMPARE, CDC, and NCBI antibiograms) as containing relevant information and therefore extracted their data. The number of entries per pathogen contributed by each database is listed in Supplementary Table S2 and summarised in Supplementary Table S3.

As the genomic component of CABBAGE entries is structured around BioSample IDs, all sequencing run IDs, sample IDs, and genome accession IDs were converted to BioSample IDs. Missing values in these genomic fields were then retrieved using BioSample IDs as the central reference. In addition, to improve the completeness of metadata fields, isolate metadata were retrieved for entries with missing values, again using BioSample IDs as the reference.

### 2.2 Data standardisation

To ensure comparability across datasets obtained from different sources, the data was mapped to a set of 27 standardised fields (Table 1), with each entry being defined by a pair of BioSample ID (genotype) and antibiotic name (phenotype). A key step in the data standardisation process involved the harmonisation of antibiotic names and their extraction from abbreviations as, for instance, a single abbreviation may refer to numerous different antibiotics across different datasets. To address this, antibiotic names were derived from the antibiotic abbreviations provided in the source dataset, referring to the paper’s full text when necessary, and standardised using the name list employed by NCBI antibiograms.

**Table 1:**
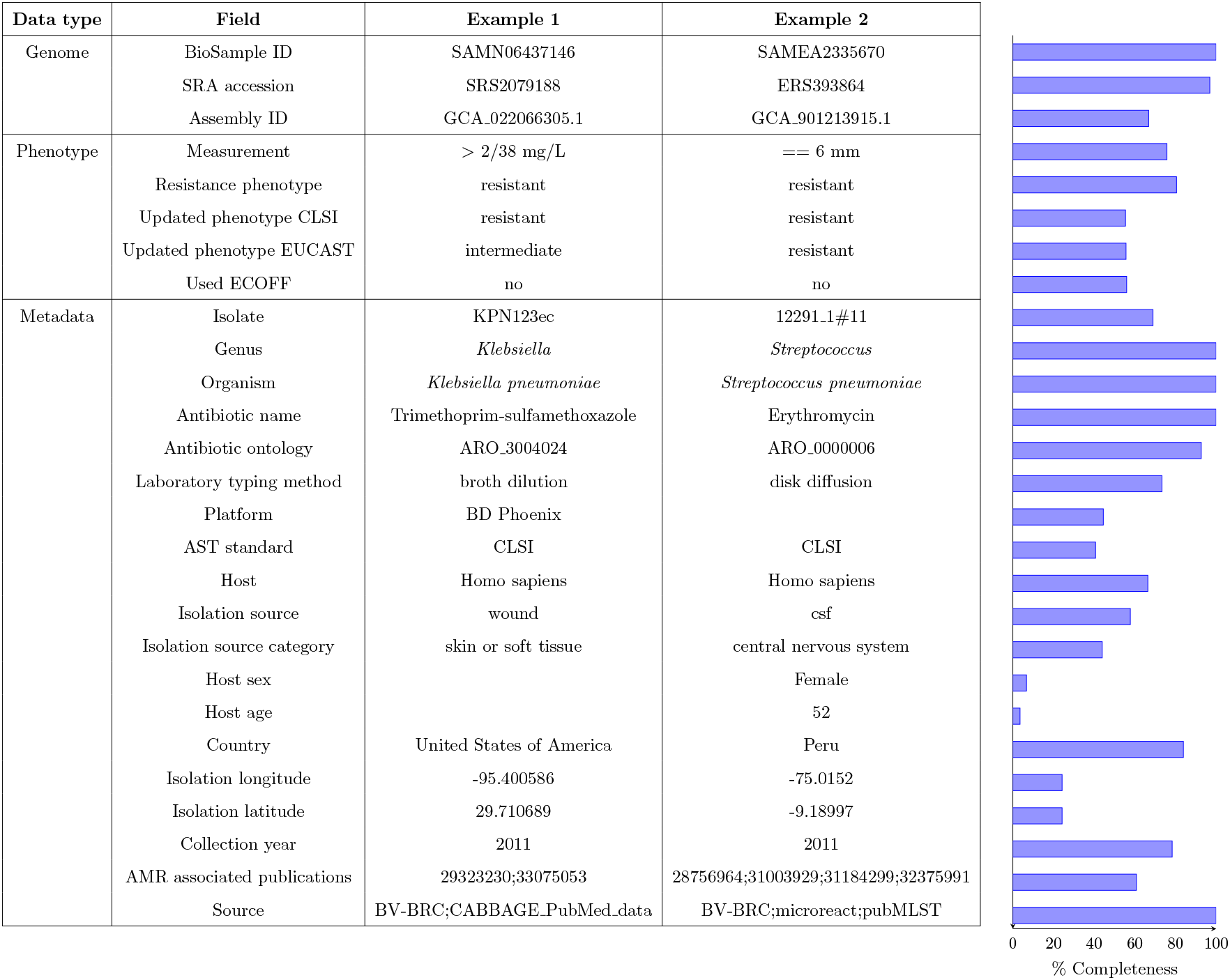
CABBAGE database fields, grouped into genomic, phenotypic and metadata categories. Two examples of CABBAGE entries are provided, as well as the percentage of completeness of each field on the right side.

Further data standardisation steps include, but are not limited to, the conversion of numerical phenotypes (e.g. 0, 1) to their corresponding classification (e.g. susceptible, resistant), the standardisation of country names based on ISO 3166-1 alpha-3 codes, standardising both broth microdilution and macrodilution to broth dilution, and standardising the host to either ‘*Homo sapiens*’ or ‘other (non-clinical isolate)’. A full description of the data standardisation process is included in the Supplementary methods (S1.5).

### 2.3 Data curation

A series of curation steps were performed to remove or correct inconsistent entries. Only those entries that were inconsistent or conflicting were manually curated. Pathogens not recorded in the WHO Bacterial Priority Pathogens List were excluded, and several species were renamed for consistency (e.g. *Salmonella typhimurium* was changed to *Salmonella enterica*). Particular attention was paid to the curation of genomic identifiers (the genomic component of CABBAGE), as well as measurement values and typing methods (the phenotypic component of CABBAGE).

#### 2.3.1 Genomic data curation

During data collection and the subsequent conversion of genomic identifiers to BioSample IDs, we identified cases where multiple genomic identifiers within a single entry did not point to the same BioSample ID (e.g., a sequencing run ID and a genome accession ID corresponding to different BioSample IDs). As the correct BioSample ID could not be determined, these entries were excluded. We also identified multiple BioSample IDs pointing to the same sequencing run ID. The corresponding entries were also excluded as these cases likely correspond to pooled isolate sequencing.

#### 2.3.2 Phenotypic data curation

Minimum inhibitory concentration (MIC) values were examined for potential errors that were subsequently removed, for example, large values inconsistent with standard doubling dilution ranges (e.g. 1,000,000). Typing methods associated with anomalous diffusion zone diameter values (e.g., small decimal values) or other irregular MIC values (e.g., series of integers outside standard dilution ranges) were manually checked against the original data sources. When errors were confirmed, both the method and the measurement unit were corrected. Original data sources were also checked to retrieve corresponding typing methods in cases where a measurement value was present without a unit and method. Further curation steps addressed anomalies in metadata fields, such as high values of host age (e.g 999) and improbable collection years (e.g. 1800) that were removed.

The final curation stage involved the de-duplication of entries, including entries that were either identical or shared the same pair BioSample ID/antibiotic name but differed in measurement value or phenotype. In the case of identical entries, duplicates were removed, and conflicting entries that could not be resolved by checking the original sources were also excluded. A full description of the data curation process is included in the Supplementary methods (S1.6).

### 2.4 Final processing

In cases where both a measurement value and categorical phenotype (e.g. resistant [R], intermediate [I] or susceptible [S]) are recorded, the phenotype is often derived using outdated or custom breakpoints. Additionally, many entries include a measurement value, but lack a corresponding phenotype. To address this, we incorporated updated phenotypes in the ‘Updated phenotype CLSI’ and ‘Updated phenotype EUCAST’ fields by applying the current 2025 breakpoint criteria from both CLSI and EUCAST, inferred using the AMR (for R) package v3.0.0 [18]. These phenotypes will be updated using the most recent breakpoint criteria in future versions of CABBAGE.

The CABBAGE database then underwent a series of consistency checks to ensure that values within each entry were consistent across fields. For instance, a testing standard can only be reported in the presence of a phenotype, a platform value should have a corresponding typing method, a measurement unit is required if a measurement value is provided alongside a typing method, and each entry must contain both a BioSample ID and either a measurement value or a categorical resistance phenotype.

### 2.5 Genome assembly and annotation

Genome annotation was conducted on genomes previously submitted to the European Nucleotide Archive and on newly assembled genomes from the AllTheBacteria collaboration [19]. Genomes were assessed using CheckM2 with 1,269 genomes filtered due to a completeness score less than 50 or contamination score greater than 5 [20]. Each genome in the CABBAGE data set was annotated using the pipeline mettannotator [21], which includes AMR annotation from AMRFinderPlus [22], providing a unified and consistent annotation set across all CABBAGE genomes.

### 2.6 Database accessibility

CABBAGE is available via the Antimicrobial Resistance Portal at EMBL-EBI (https://www.ebi.ac.uk/amr). The resource will be updated on a regular basis to incorporate new datasets and functionality. All data, including genome annotations, are made available via the EMBL-EBI FTP site (https://ftp.ebi.ac.uk/pub/databases/amr_portal) under a CC-BY 4.0 license. The data is also available in apache parquet format for raw download. Antibiograms have been deposited alongside each BioSample record. Assembled genomes are available via the European Nucleotide Archive.

## 3 RESULTS

### 3.1 Antimicrobial resistance portal

The Antimicrobial resistance portal provides a user-friendly interface to explore CABBAGE and consists of three main data resources: ‘AMR phenotypes’, consisting of antibiogram records; ‘AMR genotypes’, consisting of *in silico* annotations from AMRFinderPlus; and ‘Combined phenotypes and genotypes’, a merged dataset of phenotypes and genotypes. Within each resource, data can be filtered based on key data attributes, including species, genus, and resistance phenotype, among others (Figure 1). Once filtered, data can be downloaded in CSV format, or each individual data set can be downloaded for offline processing.

**Figure 1.**
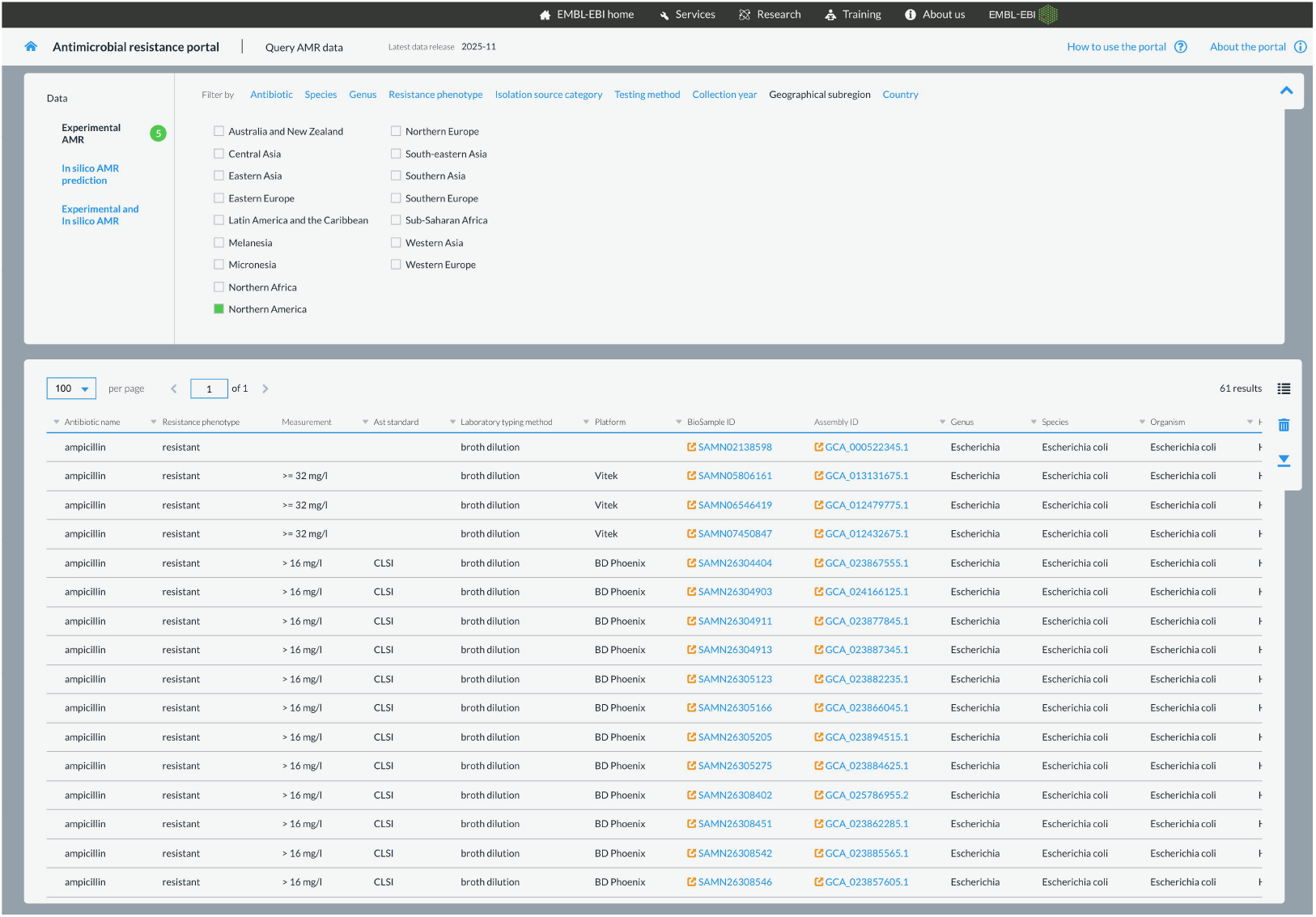
Screenshot of the Antimicrobial resistance portal, showing the CABBAGE database filtered by antibiotic, species, isolation source category and geographical region.

### 3.2 Overview of the CABBAGE database

In total, CABBAGE contains antibiotic susceptibility data and associated metadata for 170,750 sequenced isolates, with an average of 10 *±*6(SD) antibiotic susceptibility test results per isolate. The database contains *>*1.7 million entries, each entry being defined by a BioSample-antibiotic combination with an associated AST result. The AST result can be represented by a measurement value and/or a categorical phenotype (e.g. S, I or R). Both a measurement value and a categorical phenotype are recorded for *>*964,000 entries, and both a measurement value and an updated phenotype, which we derived by comparing the measurement value to the latest CLSI and EUCAST breakpoints, are recorded for *>*1.1 million entries (Table S5). Table 1 summarises field completeness alongside two examples of CABBAGE entries. While the fields defining entries are 100% complete, some metadata fields, such as host sex and age, are sparsely populated but are retained in the anticipation that coverage may improve as the database expands.

The number of entries obtained from, and shared between, different data sources is summarised in Figure 2a. BV-BRC is the largest single data source, contributing just over 764,000 entries, followed by datasets derived from our literature review (89 publications), with over 560,000 entries. We observed high redundancy between data sources, which allowed us to identify and remove problematic entries during the database deduplication process. BV-BRC and the data sourced from publications have the largest number of entries in common with just each other (*>*142,000), followed by NCBI antibiograms and NDARO (*>*126,000).

**Figure 2.**
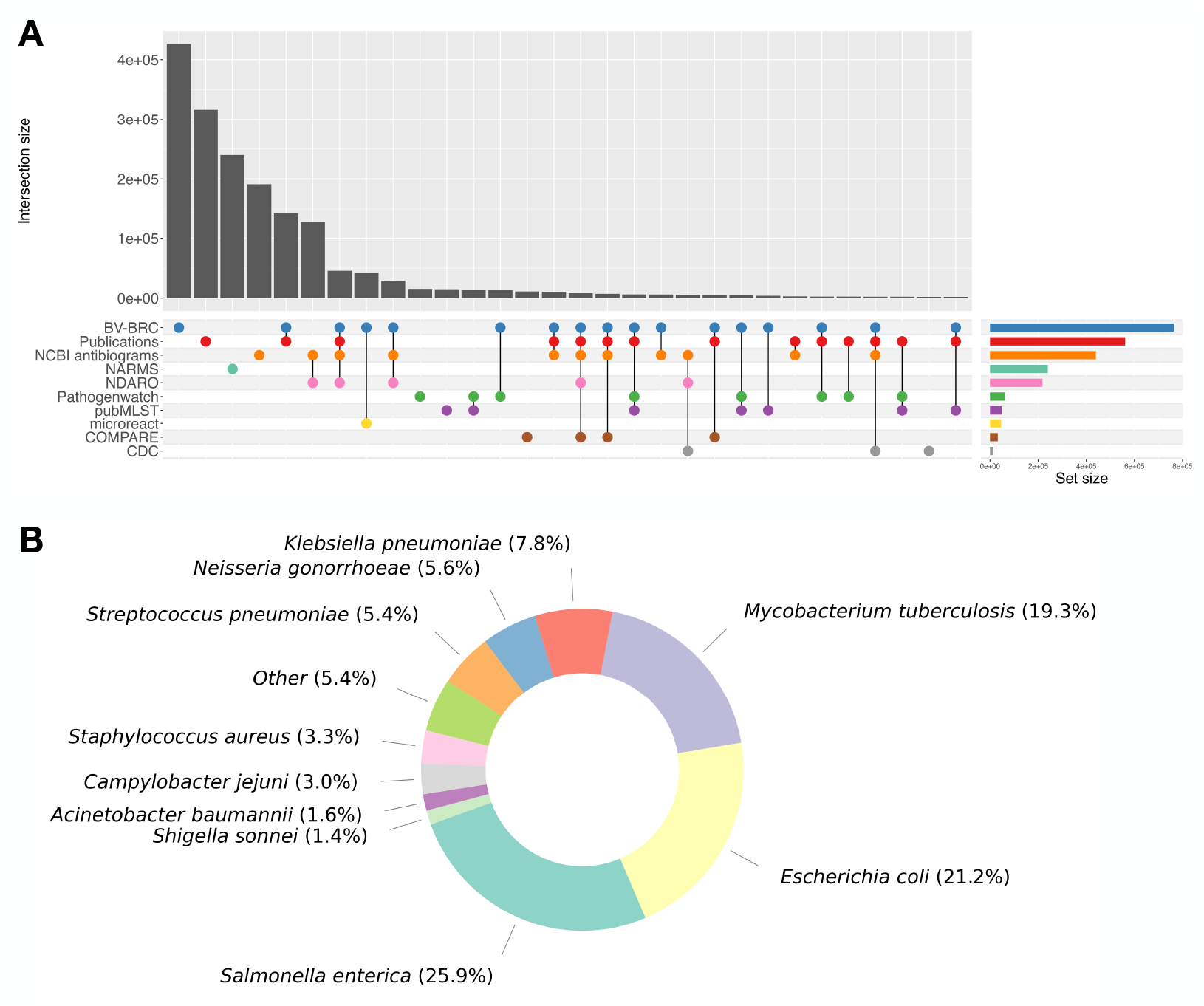
A) Sources of CABBAGE data, in numbers of entries. Only sets with a minimum intersection size of 2,000 are displayed. The full UpSet plot is provided in Supplementary Figure S1. B) Distribution of the 10 most common pathogens in CABBAGE. All remaining species are grouped under ‘Other’. Percentages represent the number of entries (pathogen-antibiotic pairs).

Finally, the data is not evenly distributed across pathogens, with three species representing more than half of entries: *Salmonella enterica* (25.9%), *Escherichia coli* (21.2%), and *Mycobacterium tuberculosis* (19.3%) (Figure 2b). In contrast, other pathogens are under-represented, with some species being represented by a single unique isolate: *Enterobacter cancerogenus, Serratia fonticola, Enterobacter mori, Enterobacter soli, Serratia nematodiphila, Providencia alcalifaciens, Proteus alimentorum* and *Enterobacter sichuanensis*.

### 3.3 Spatio-temporal distribution of CABBAGE data

As expected, we observed substantial geographical disparities in the AMR data, with some regions, such as Central Africa and Central America, being sparsely sampled (Figure 3a). The US has the largest data coverage (*>*698,000 entries), in part due to their formidable AMR surveillance programs and associated databases (NARMS, NDARO and CDC). Additionally, these inconsistencies in global data distribution may reflect differences in how readily organisations make data available following collection. Variations are also seen in the frequency of resistance phenotypes between different countries, as illustrated in Figure S4. However, caution should be taken when comparing resistance frequencies between countries due to varying levels of data coverage and ascertainment bias due to the fact that not all isolates are sequenced, and in some settings, sequencing is prioritised for those isolates that are phenotypically resistant to at least one antibiotic [23].

**Figure 3.**
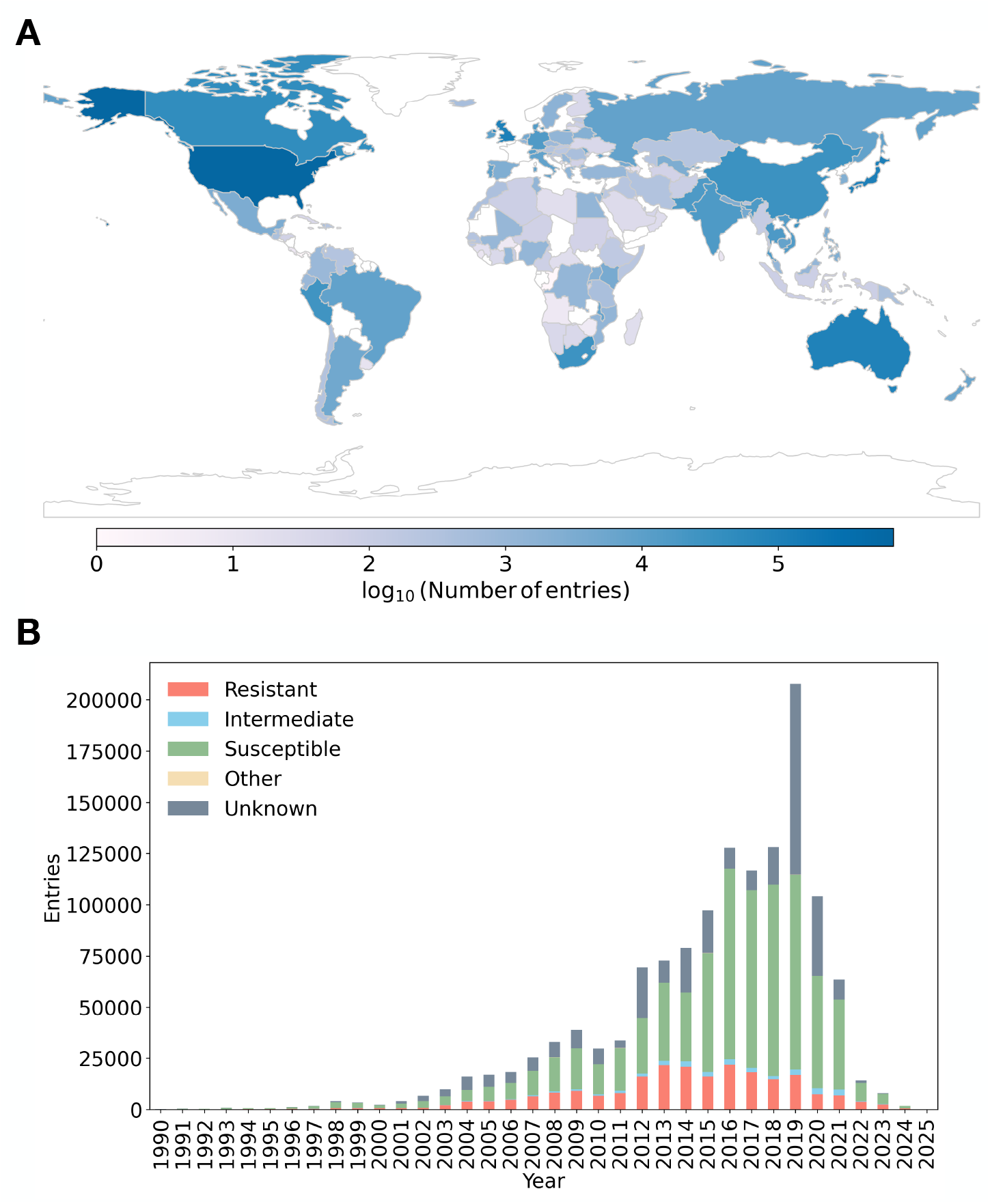
A) Geographical distribution of entries (log_10_). Countries that are not represented in CABBAGE are shown in white. B) Distribution of CABBAGE entries per collection years. The category ‘other’ includes susceptible-dose dependent and non-susceptible. Frequencies of each categorical phenotype per year are shown in Supplementary Figure S5b.

New AMR datasets are being published at an increasingly rapid pace, as illustrated in Figure 3b. There is a marked increase in the number of entries from approximately the year 2000 onwards, likely reflecting advances in WGS, AMR surveillance and data sharing. However, we also observed a sharp decline in the number of CABBAGE entries corresponding to recent years. This decline may be purely technical, reflecting a lag in the incorporation of recent data into public databases. Alternatively, it could indicate a genuine reduction in global AMR data collection, resulting from the substantial impact of the COVID-19 pandemic on AMR surveillance programs and academic research [24, 25, 26]. Future updates of the CABBAGE database incorporating more recent data sources will help disentangle the relative contributions of these two, not mutually exclusive, hypotheses.

### 3.4 Antibiotic resistance frequency in CABBAGE

The percentage of resistant isolates varies widely between drugs (Figure 4a). Among antibiotics with the greatest data coverage (at least 10,000 isolates), ampicillin showed the highest resistance frequency (40%), whereas bedaquiline had the lowest (1.2%). These differences likely reflect both the time since each drugs introduction and their overall levels of clinical use. However, the CABBAGE dataset as a whole is expected to be biased toward resistant phenotypes (22% of CABBAGE entries with a categorical phenotype are resistant), since such isolates are more often sequenced and reported.

**Figure 4.**
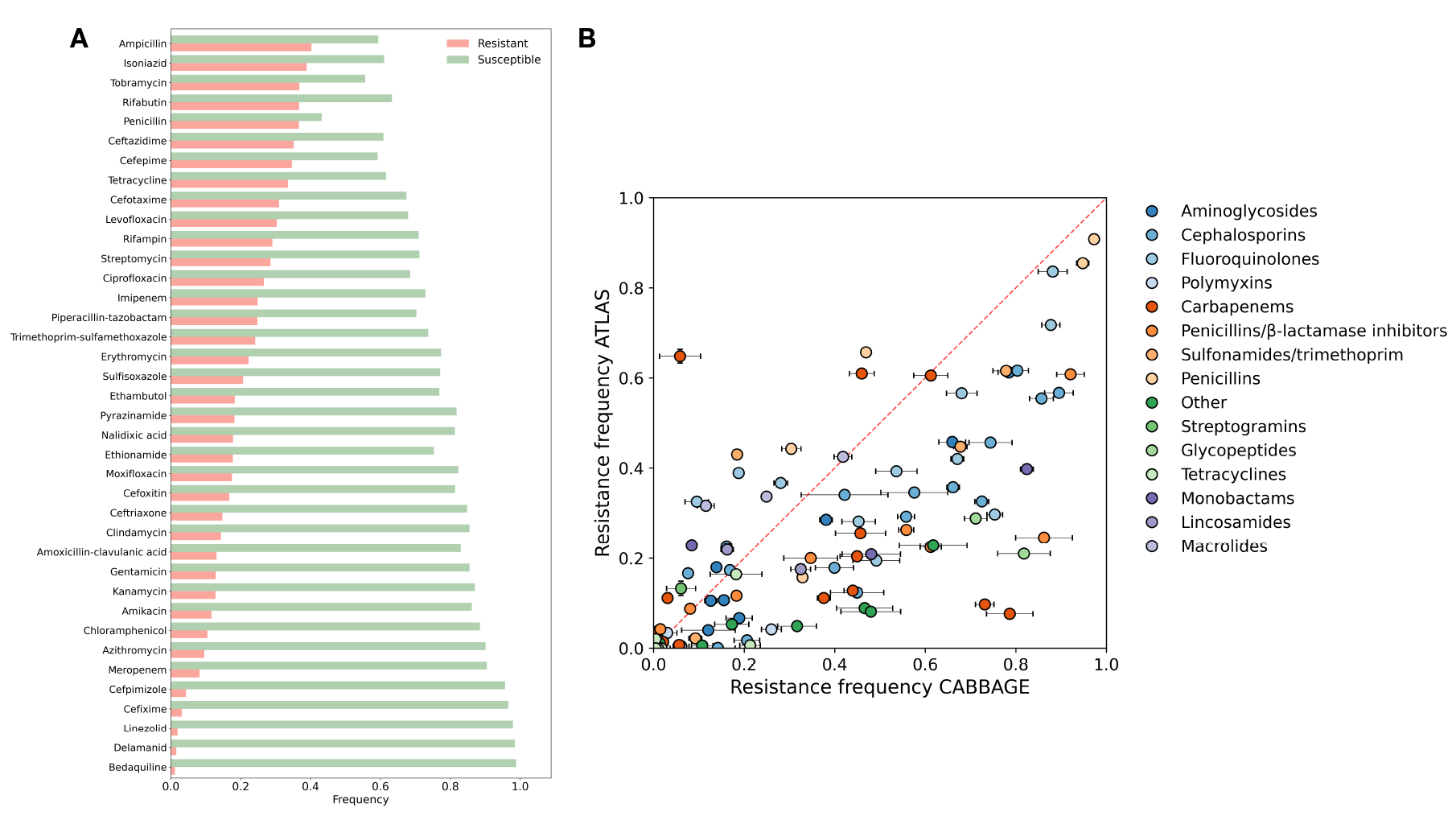
A) Frequencies of resistant (red bars) versus susceptible (green bars) phenotypes across the CABBAGE dataset, by antibiotic. We used here the phenotypes extracted from the original data source, not the updated phenotypes, and the data was restricted to antibiotics with at least 10,000 entries in CABBAGE. Absolute numbers of susceptible and resistant entries are displayed in Supplementary figure S7. B) Frequencies of resistance in CABBAGE versus ATLAS. Each data point represents a unique pathogen-antibiotic pair present in both datasets, restricted to data collected between 2004-2023 to reflect the range of collection dates in ATLAS. A scatter plot without this date restriction is presented in Supplementary Figure S8. Pathogen-antibiotic pairs with at least 100 entries in both CABBAGE and ATLAS are shown. Error bars represent 95% CI and colours are representative of different antibiotic classes. The CABBAGE phenotypes presented here represent the phenotypes derived from the original datasets, rather than the updated phenotypes.

To evaluate this potential bias, we compared the resistance frequencies of pathogen-antibiotic pairs shared between CABBAGE and the ATLAS database [17] (104 pairs; Figure 4b). ATLAS serves as a useful comparator because it compiles results from large global surveillance studies (*>*900,000 isolates). Logistic regression analysis revealed that resistance was significantly more frequent in CABBAGE than in ATLAS (OR = 1.6, *p <* 0.001). Nonetheless, this trend was not universal: 66 pathogen-antibiotic pairs showed significantly higher resistance in CABBAGE, while 29 showed greater susceptibility (Fishers exact test, FDR adjusted *p <* 0.05).

## 4 DISCUSSION

Comprehensive, large-scale datasets integrating bacterial genome sequences with associated phenotypic antibiotic susceptibility data are essential for the development of genotype-to-phenotype predictive models to further our understanding of AMR mechanisms and to inform the design of improved diagnostics. Although a number of databases provide such information, up until now this data has not been collated and processed into a single uniform format, limiting its collective analysis. A substantial volume of data also remains in the public domain across multiple publications, with more being added on a weekly basis. To address this challenge, we developed CABBAGE, a comprehensive, global, curated genotype-phenotype AMR database for WHO Bacterial Priority Pathogens. CABBAGE is the largest database of its kind, freely available via the Antimicrobial Resistance Portal at EMBL-EBI (https://www.ebi.ac.uk/amr), facilitating user-friendly access.

Through an extensive literature review and examination of public databases, we extracted data on *>*170,000 unique sequenced isolates with associated phenotypic AMR data, totalling *>*1.7 million genome-phenotype pairs. In addition, each entry is associated with metadata where available, such as the isolate collection date, isolation source, and geographic location. The data was standardised into a uniform format and curated, where necessary, to improve consistency, accuracy, and completeness. CABBAGE constitutes a wealth of information that facilitates benchmarks of genotype-to-phenotype predictive models, with the potential to enhance our understanding of the genetic mechanisms of AMR, including the identification of novel resistance determinants. Additionally, CABBAGE enables the analysis of spatiotemporal resistance trends for AMR surveillance, and can be used to enhance clinical diagnostic tools, such as genome-based antimicrobial susceptibility testing.

Since the preprocessing pipeline may influence the downstream inference [27, 28], we recommend that data producers make genotype data available in raw format, namely as sequence reads submitted to the ENA. Additionally, antibiogram data should ideally be submitted to BioSamples. As part of this work, we have created new guides to help data generators archive their data: https://www.ebi.ac.uk/amr/amr_submission_guide/. By submitting both genotype and phenotype data to biomedical data archives, this achieves longevity of storage in a uniform format, maximising reusability, and avoiding being tied to a single research project. Prediction methods would likely need to first pre-process raw read data into genomic features, which is important to do in a consistent manner across the dataset. Assemblies, gene calls or variant calls agreed by the community are a useful resource for reusers of this data, and our consistent assembly and annotation of data was designed to save computational time and improve consistency for downstream users of CABBAGE.

The data within CABBAGE exhibits multiple layers of heterogeneity, for example, phenotypes vary from quantitative MICs and disk diffusion data obtained from a variety of AST methods, to S/I/R classifications obtained using guidelines that may be outdated. We addressed this by providing additional updated phenotypes based on current EUCAST and CLSI guidelines. The species in CABBAGE are also unevenly distributed, with *S*.*enterica* alone accounting for just over 25% of all entries, and at the other extreme, a number of species are only represented by a single isolate. Similar heterogeneities are found for the antibiotics, publications, countries represented, and the years of collection. We specifically highlight a lag in the availability of new datasets over recent years, and hypothesise that this could be due to the impact of the COVID-19 pandemic on AMR research, a delay in the incorporation of new data into databases, or a combination of both these factors.

Lastly, we note that the fraction of resistant isolates is likely to be over-estimated, as many groups tend to perform AST on all isolates of interest, but only sequence those that are resistant. We illustrate this bias towards resistance through a comparative analysis of CABBAGE against the AMR dataset ATLAS, showing that, overall, there is a greater likelihood of resistance in CABBAGE relative to ATLAS. It is possible that CABBAGE could be used to systematically identify gaps in surveillance, even though, depending on the sampling methodology and the resources available, such gaps may or may not be easy to fill. Future updates to CABBAGE will focus on extending the database, particularly for under-represented species, and applying the most recent WHO Bacterial Priority Pathogen List (2024).

In summary, CABBAGE consolidates data from existing resources into a single, unified database. Through the integration, standardisation and curation of information that was previously fragmented, CABBAGE provides a strong foundation to advance research and innovation in the AMR space. Additionally, it highlights the need to adopt a standard format for sharing genotype-phenotype data [29], which will facilitate, and aid the automation of, its extension as additional data is generated.

## 6 ACKNOWLEDGMENTS

This work is supported by funding from the UK Medical Research Council (MR/Z505547/1). LC is supported in part by the NIHR Imperial Biomedical Research Centre (BRC). RD, ED and LC acknowledge funding from the MRC Centre for Global Infectious Disease Analysis (reference MR/X020258/1), funded by the UK Medical Research Council (MRC). This UK funded award is carried out in the frame of the Global Health EDCTP3 Joint Undertaking. AA, TG, BEH, JK, SO, OK, NMR, AS, AW, GY, ADY, RF and JL acknowledge funding from Wellcome Trust (228142/Z/23/Z) and European Molecular Biology Laboratory. The authors would like to acknowledge Antonio Pedrotta, Rupal Ramesh Shah, Atharv Naik, Dipayan Gupta, Zahra Waheed, Lucy Brooks, Jorge Batista da Rocha, Kamalkumar Dodiya, Stefano Giorgetti, Tim Dallman, Elita Jauneikaite, Nicholas Croucher, Yonatan Grad, Catrin Moore, Gus Hamilton, Gwen Knight, Till Bachmann, Edward Feil, Sarah Fortune, Xavier Didelot, Sean Nachtrab, the PHA4GE data structures working group, the ESGEM-AMR working group and the B2B2B AMR diagnostics network for their ongoing support of the project. The authors are grateful to all the maintainers of and contributors to all the databases, as well as the authors of all the publications used in this work. For the purpose of open access, the author has applied a CC BY public copyright licence to any Author Accepted Manuscript version arising from this submission.

