## Supplementary Files for "A comprehensive AMR genotype-phenotype database (CABBAGE)"

### S1 SUPPLEMENTARY METHODS

#### S1.1 Literature review and data extraction

A literature review was carried out, with the initial aim of collecting publicly available datasets containing at least 100 sequenced isolates of bacteria on the WHO's Priority Pathogens List, with antimicrobial resistance phenotypes available for at least one drug. The process recommended by Bramer *et al.* [1] was followed, with the use of the Embase database as the primary source of information. During the refinement process, the minimum sample size for *M. tuberculosis* was increased to 200 due to the large number of available datasets, and decreased to 50 for less commonly represented bacterial pathogens.

A total of six search terms were used, as summarised in Supplementary Table S1. Where a search term generalises a previous one, we assumed that all previous hits would be included, and thus compute and report those statistics only on the new hits. All search terms were restricted to English-language papers, and this included papers from 2010 until the date of the search (July 2022). The search identified 228 unique papers among the 4,325 considered, for an overall precision of 5%. In total, 232 datasets were collected, with a dataset defined as a collection of isolates belonging to a single species, as some of the papers included more than one priority pathogen bacterial species. A further non-systematically identified sample of 28 papers was added, bringing the total number of papers to 256.

During the literature review, papers were excluded if they fell into certain categories. The main categories of excluded results were as follows:

- off-topic papers, such as papers on viruses, fungi, malaria or cancer;
- review papers, position papers, and conference abstracts;
- papers whose sample size was too small (e.g. case studies);
- epidemiology papers (e.g. prevalence papers);
- laboratory or functional genomics papers;
- antibiotic, phage, or antimicrobial peptide identification or design papers;
- studies doing selective sequencing (e.g. known resistance-associated genes).

A number of papers had their data sourced from previously published studies. Any such paper was included provided that it explicitly made the aggregated genotype and phenotype data available, rather than merely referencing the previous studies; otherwise, the paper was not included, while the referenced previous studies were individually considered for inclusion if they met all the search criteria.

Among the papers identified through the search, eight bacterial pathogens were represented only once, namely, *Escherichia albertii*, *Enterobacter cloacae*, *Enterococcus faecalis*, *Haemophilus influenzae*, *Neisseria meningitidis*, *Serratia marcescens*, *Staphylococcus pseudintermedius*, and *Vibrio cholerae*. Of those, only *H. influenzae* and *S. marcescens* are WHO Priority Pathogens, although several others have appeared on the US CDC [2] or India's Priority Pathogen Lists [3]. The six non-priority pathogens were excluded from analysis, leaving 248 datasets from 224 papers.

For each dataset, an attempt was made to extract the genotype accession numbers as well as the phenotype tables, in the first instance from the Supplementary Materials of the paper. If this was not successful, the full text of the paper was analysed to identify included accession numbers (by textual search for the terms "access", "avail", and "depos") and phenotype tables (by visual inspection of the tables and figures). As a result of one of the types of data (either genotypes or phenotypes) being unavailable, the original set of papers was narrowed down to 127 datasets from 107 papers.

Papers reporting the genotypic prediction of resistance and its concordance with phenotypic resistance were excluded if they did not explicitly specify the discrepant isolates, or if they did but only provided the concordant phenotypes implicitly as the results of a prediction tool, since such tools tend to be updated over time and may yield predictions that disagree with their original ones. The sample size was also reduced for several datasets, as not all strains under study were publicly available, and 14 were then excluded due to the remaining sample size being too small (below 100), resulting in 113 datasets from 94 papers. As some datasets used identical sources for both their genotypic and phenotypic data, they were deduplicated, leaving 110 unique datasets from 91 papers.

##### S1.2 Initial data processing

Several papers only had their data available in formats that are challenging to process via automated scripts, such as PDF files containing tables or PNG figures. In this instance, the paper's corresponding author was contacted with a request to provide the data in a machine-readable format. The authors were contacted a second time after 5 working days, and if no reply was received after a further 5 working days, the data was considered unusable for data collection and removed from further consideration. Seven out of eleven corresponding authors complied with the request within 10 working days. Four failed to respond, resulting in an attempt to "manually" convert 3 PDF files into CSV files. One dataset whose author did not respond to the request was excluded due to only having phenotype data in the format of a phylogenetic tree with no labels. No contact was made with authors of papers that did not include both genotype and phenotype data, even if these were stated to be available upon request.

For each dataset with publicly available data, we extracted tables containing the phenotypes as well as the accession numbers for the genotypes, with a preference for raw sequencing data when it was available; we used assembled contigs when no raw sequencing data was available, but did not include any papers that only provided access to variant calls as these may be difficult to reproduce. The hyperlink corresponding to each table and study was recorded. When the article was freely available in PMC we used its PMC version as the source of the data; however, some of the articles were behind a paywall and the links we provide to the data in those may not be accessible at all institutions, hindering the open access of the data and the reproducibility of our results.

The tables were first standardised into a CSV format (comma-separated rather than semicolon-separated) manually due to the variability of formats. For Excel files, rows before the header and after the final row were removed; sometimes this necessitated the removal of columns (for instance, if the header was on two levels, with the first level providing the drug name and the second level stating if the reported result was experimentally measured or computationally predicted, the computational predictions had to be removed to ensure lossless conversion into a CSV format). For Excel files with multiple sheets, only the relevant tabs were converted, and when multiple datasets were provided as separate sheets, each sheet was treated separately and then merged if appropriate (e.g. a training dataset and a testing dataset could be merged into a single overall dataset if their formats were compatible).

For those genotypes available only as an INSDC project, the RunSelector tool (NCBI) was used to extract the correspondence between internal IDs/sample names and accession numbers. For those papers where the genotype and the phenotype information were provided as separate tables (including those with genotypes available via an INSDC project), the tables were merged using an inner join (i.e. keeping only those isolates with both phenotype and genotype information available), using manually determined identifier columns in non-trivial cases, except when the manuscript clearly specified a default phenotype that was then assumed for all samples present in the genotypic table but absent in the phenotypic table. One additional dataset was excluded due to having no identifiers usable for matching and no reply to a request to its authors for a dictionary to establish the correspondence. This left a total of 108 datasets from 89 papers.

##### S1.3 Database analysis

The following lists the URL's for databases that were assessed for integration into CABBAGE:

- Bacterial and Viral Bioinformatics Resource Center (BV-BRC), at https://www.bv-brc.org;
- EnteroBase, at http://enterobase.warwick.ac.uk;
- MGTdb, at http://mgtdb.unsw.edu.au;
- Microreact, at http://microreact.org;
- National Antimicrobial Resistance Monitoring System for Enteric Bacteria (NARMS), at http://www.cdc.gov/narms/index.html;
- National Database of Antibiotic Resistant Organisms (NDARO), at http://ncbi.nlm.nih.gov/pathogens;
- PathogenWatch, at https://pathogen.watch;
- pubMLST, at http://pubmlst.org.

Unfortunately, EnteroBase does not contain any AMR phenotype information except for a small number of *Helicobacter pylori* isolates, while MGTdb only contains such information for *Bordetella pertussis*, where it is computationally rather than experimentally predicted. We thus focus our initial efforts on the remaining 6 databases: BV-BRC, microreact, NARMS, NDARO, PathogenWatch, and pubMLST, as well as the European COMPARE Consortium.

###### S1.3.1 BV-BRC

For the BV-BRC database (formerly known as PATRIC) we followed the recommendation of Van Oeffelen *et al.* [4] and downloaded the entire contents of the database from the links at https://docs.patricbrc.org/user_guides/ftp.html. Because many phenotype entries lacked a genomic identifier and included only a sample name, the corresponding sequencing run IDs were retrieved either from the source publications (2,362 isolates; PubMed IDs 32025709, 30373719, 31537784, 29133554, and 34485958) or from the BioSample database, when the search returned a single matching BioSample ID whose species matched that of the corresponding BV-BRC entry (3,676 isolates). Filtering the BV-BRC entries to species on the WHO Priority Pathogen list resulted in a total of 83,267 unique isolates. The final number of entries sourced from BV-BRC after processing are summarised in Table S2.

###### S1.3.2 The European COMPARE Consortium

The European COMPARE Consortium [5] has previously attempted to standardise the reporting of AMR phenotypes, and there are a total of 2,570 isolates (790 *Escherichia coli* and 1,780 *Salmonella enterica*), together with antibiograms, included in its AMR Data Hub. We include this data from the AMR Data Hub in the current analysis because, although much of it is also found in other sources, the formatting of the antibiogram data allowed for straightforward integration into existing workflows. However, we note that for one of the 12 studies included in the AMR Data Hub (PRJDB7087), each of the isolates has a repeated value for three antibiotics (aztreonam, cefotaxime, and cefotiam), so these antibiotics are excluded from our analysis in this study.

###### S1.3.3 Microreact

Since Microreact projects are not explicitly linked to from its homepage, except for the Showcase selection [6], we followed each of the five showcased datasets focused on bacterial pathogens in the WHO Priority Pathogens List to its original publication. The first was already included in our search [7]; another [8], although not identified in our search, lacked antimicrobial resistance data in the Microreact table, which was instead extracted from the original publication. The remaining three [9, 10, 11] either did not contain any antimicrobial resistance data, or had fewer than 100 strains, and thus were not used. We also examined the 21 Microreact collections at `https://www.microbiologyresearch.org/content/microreact` to identify any that had additional data meeting our search criteria, and found five matching datasets, one of which was included in our original search.

###### S1.3.4 NARMS

NARMS primarily contains information on foodborne pathogens. We query it for genotyped isolates from 1999 onwards with AST-based phenotypes, excluding *Vibrio* which is not part of our search targets.

###### S1.3.5 NDARO

NDARO contains information from NCBI on a number of pathogens and their antimicrobial resistance. It can be accessed via FTP at `ftp://ftp.ncbi.nlm.nih.gov/pathogen`. Each genus or species of pathogen is collected into its own folder, and the AMR subfolder contained in each of those provides a file that specifies the combination of accession number and AST profile, along with many other metadata fields. We downloaded these files for each of the bacterial pathogens on the WHO Pathogen Priority list.

###### S1.3.6 PathogenWatch

The PathogenWatch database is organised into 89 collections of isolates. Of those, 44 have fewer than 100 isolates, and are therefore excluded from our analysis. Of the remaining 45 collections, a further 3 contain pathogens not targeted by our search, and one (for the EuroGASP 2013 collection) contains no identifiers that could lead to accession numbers. The 41 remaining collections were processed by downloading the metadata and the AMR profile in csv format, merging them using a custom script and filtering out any isolates which only had the genotype or the phenotype available, but not both, as well as querying EBI for obtaining run accession numbers from the sample accession numbers when needed. Finally, the files were processed to remove duplications.

###### S1.3.7 pubMLST

Since pubMLST is primarily focused on providing multilocus sequence type information, the incorporation of resistance profiles seems to be a secondary goal. For this reason, most of the pathogens of interest to us did not contain enough isolates with whole genomes as well as resistance profiles to be selected, with the following exceptions (Table S2), found by querying the genome collection for each relevant species with the 'Field NOT null' search specification, where Field was 'resistance profile', 'drug resistance', or antibiotic names combined via OR.

##### S1.4 Metadata extraction

To improve the completeness of the database, relevant metadata was extracted directly from the manuscripts and supplementary materials of the 108 datasets collected from 89 papers during the literature review. This metadata was recorded in a metadata CSV file, separate to the CABBAGE dataset, with the aim of incorporating metadata present in the manuscript or supplementary materials, but absent in the original datafile. Each manuscript and its associated supplementary materials were manually examined and, where relevant information was identified, it was highlighted within the text and recorded within fields based on those used by NCBI BioSample Antibiograms. These fields include 'Laboratory typing method', 'Laboratory typing platform', 'Testing standard', 'Testing standard documentation', 'Vendor' and 'Laboratory typing method version or reagent'. In addition to these fields, we defined three additional fields: 'Temperature ( °C)', 'Incubation time (hrs)' and 'Conditions'. However, due to the sparsity of information available for these three fields (the temperature, incubation time and conditions were only specified in 6,

7 and 2 papers respectively), this information was not incorporated into the final database.

For a number of papers, the methods and/or testing standards were provided for specific antibiotics, and so these were listed alongside the corresponding antibiotic in the field 'Laboratory typing platform - antibiotic specific' or 'Testing standard - antibiotic specific'. Where different methods were applied to different sets of isolates within a paper, the entry was split, and this was indicated in the 'Duplicate' field with a 'yes' or 'no' entry. An additional 'Notes' field includes any information relevant to the metadata, for instance the specific breakpoints that were used, if different methods were used for different subsets of isolates, and if these subsets were ambiguous. Note that fields such as 'Notes' and 'Duplicate' are not included in CABBAGE, as their purpose was to provide additional information required for the decision-making process to incorporate metadata into CABBAGE.

##### S1.5 Data processing and standardisation

Bacterial pathogens not included in the WHO Priority Pathogen list were removed from further analysis, and several species were renamed for consistency as follows: *Salmonella typhimurium*, *Salmonella Senftenberg* and *Salmonella enterica subsp. enterica serovar Kentucky* to *Salmonella enterica*; *Enterobacter cloacae complex* and *Enterobacter cloacae group* to *Enterobacter cloacae*; and finally *Providencia* species to *Providencia stuartii.*

It was noted that a number of PathogenWatch datasets were derived from the 'AMR profile', consisting of phenotypic resistance predictions based on antibiotic resistance genes. As a result, these datasets were removed from further analysis, reducing the number of datasets from PathogenWatch from 41 to 21. Two additional datasets [12, 13] derived from the literature search were excluded from further analysis as an excessive number of assumptions would otherwise be required to reliably interpret the phenotypic data.

Antibiotic abbreviations were derived from the antibiotic name provided in the original dataset, if available, based on the names and corresponding abbreviations used in the AMR package [14]. To ensure accuracy in the assignment of antibiotic names in cases where only the abbreviation is provided in the original dataset, the manuscript and supplementary materials were referred to. In some cases a single antibiotic abbreviation corresponds to multiple antibiotic names across different data sources, for instance, the abbreviation CFX was used to denote either cefuroxime, cefixime or ceftriaxone. The abbreviations were therefore converted into CXM, CFM and CRO respectively. Additionally, several antibiotic names were standardised to align with those given in the AMR package, for example co-amoxyclav was changed to amoxicillin-clavulanic acid and co-trimoxazole to trimethoprim-sulfamethoxazole. Entries with only the antibiotic class recorded were not incorporated into CABBAGE if the specific antibiotic could not be determined from the original paper or Supplementary Materials.

In several instances, commonly used antibiotic abbreviations in the source materials conflicted with the abbreviation scheme used here. This was particularly apparent for tetracycline and streptomycin that frequently have the abbreviations TET and STR respectively, referring to tetroxoprim and streptoduocin in the AMR package. These discrepancies were resolved through manual verification against the antibiotic names provided in the source materials.

A variety of phenotypes are recorded in CABBAGE, and they are as follows: susceptible, resistant, intermediate, susceptible-dose dependent and non-susceptible. Where possible, phenotypes were standardised, for example, ‘S’ and ‘sensitive’ were changed to ‘susceptible’ and ‘R’ to ‘resistant’. Several datasets represent phenotypes numerically, for instance, 0 may represent susceptible and 1 resistant. In these cases, the original source was consulted to convert these numbers to their corresponding R/I/S classification.

The testing standard refers to internationally recognised guidelines, such as those set out by CLSI, which define the interpretive criteria (breakpoints) for classifying antibiotic susceptibility. For consistency, the full organisation names were converted into their corresponding abbreviation, for example, ‘Clinical and Laboratory Standards Institute’ was changed to CLSI. Ambiguous testing standards were omitted, for instance if multiple standards were reported for a single entry.

Further steps include the standardisation of the host information to either ‘*Homo sapiens*’ or ‘other (non-clinical isolate)’, with the latter encompassing all non-human entries such as animals, the environment, and food. The isolation countries were standardised using ISO 3166-1 alpha-3 codes (ISO3), which are three-letter country codes defined by the International Organisation for Standardisation (ISO). Several entries could not be converted into an ISO3 code because they do not correspond to a single country, for example, 'Caribbean’. Several country names were also updated to reflect current official naming conventions, for instance, Ivory Coast is officially designated as Côte d'Ivoire under the ISO standard, and corresponds to the ISO3 code CIV.

##### S1.6 Data curation

MICs are recorded in various formats, including integers (e.g. 2), floats (e.g 0.5), intervals (e.g. 2,5) and combinations (e.g. 2/38). It should be noted that MICs for antibiotic combinations (such as trimethoprim-sulfamethoxazole) are given as either a single MIC, or a MIC combination listing the concentrations of each component separately, depending on the format given in the original datafile. Several MICs were excluded due to probable reporting errors. This includes unusually large values inconsistent with standard doubling dilutions (e.g. 42,023 and 1,000,000), zero or null values (0.0 or 0) reported alongside 'broth dilution' as the laboratory typing method, incomplete MICs (e.g. .016), incomplete antibiotic combinations (e.g. 0.06/, /64, /128), and values with other formatting anomalies such as trailing full stops (e.g. 0.5/9.). Where possible, the measurement sign associated with each MIC was collected directly from the dataset, or otherwise recorded as '=='. MIC intervals represent the range of MIC values that lie between two defined thresholds, and so are assigned the measurement sign '>,<'. Several categorical phenotypes were removed from the database (decreased susceptibility, high level resistance, SYN-R and SYN-S), while the following categorical phenotypes were retained : susceptible, resistant, intermediate, non-susceptible, and susceptible-dose dependent.

Two measurement units are reported in CABBAGE (mg/L and mm), however if no measurement unit is provided in the original dataset, it was derived from the AST method where possible. For example, if the laboratory typing method is broth dilution, E-test or agar dilution, the measurement unit is recorded as 'mg/L', whereas 'mm' is used in the case of disk diffusion. Mismatches between the measurement unit and laboratory typing method were identified, such as entries with 'mg/L' recorded alongside 'disk diffusion'. One dataset [15] provides the units 'mg/ml' for *Mycobacterium tuberculosis* and ethionamide, however this was presumed to be an error due to the numerical values involved and changed to 'mg/L'.

The laboratory typing method field refers to the AST methodology used to generate the MIC and/or resistance phenotype, with entries recorded as either broth dilution, E-test, agar dilution or disk diffusion. It should be noted that within CABBAGE, the term 'broth dilution' is used to represent both broth microdilution and macrodilution. Where entries lacked a laboratory typing method, the metadata file (as described in Supplementary section S1.4) was consulted and the method was only recorded if it was unambiguous, for example, if a single method was used across all isolates within the study. If entries lacked a laboratory typing method that could not be resolved via this strategy, it was derived from the laboratory typing platform where possible, for example, if the laboratory typing platform was recorded as Vitek, Sensititre, BD Phoenix, Microscan or MGIT900, the method was recorded as broth dilution.

Entries for age were rounded to the nearest whole number, and those that appeared to be a reporting error, such as 999, were removed. For consistency between entries with different date formats, only the collection year was recorded rather than the full date. The only collection year to be removed was '1800', as this was believed to be a reporting error.

AMR associated publications correspond to specific publications indexed in PubMed, enabling users of CABBAGE to refer to the original data source. However, database-derived PubMed IDs should be interpreted with caution, as they don't consistently correspond to the correct publications. Several entries are associated with multiple PubMed IDs and are recorded as such, whilst other entries have no associated PubMed ID, for example if the genome and antibiogram data were deposited directly to NCBI without a corresponding publication.

The final stage of curation involved removing duplicate entries by grouping together records sharing the same BioSample ID and antibiotic name. This step also helped identify entries with conflicting phenotypes. A conflicting phenotype was defined as differing measurement values obtained using the same AST method, or - when no measurement was available - differing SIR phenotypes derived from the same method. Groups containing conflicts were discarded, except when the conflicting entries originated from the same source, in which case within-study replicates were retained. Groups without conflicts were further deduplicated as follows: entries were first ranked by the number of filled phenotype fields (see Table 1), giving highest priority to measurement values (for example, an entry with only a MIC value was preferred over one containing only SIR phenotype, AST method, and platform). The entry with the highest score was selected as the representative of the group, except when conflicting entries originated from different AST methods, in which case both entries were retained as representatives. Finally, a majority-rule consensus was applied to metadata fields, and the resulting consensus values were combined with the phenotype fields of the representative entry or entries.

To address the use of outdated interpretive breakpoints to infer phenotypes, we incorporated updated phenotypes, applying the current breakpoint criteria from both CLSI and EUCAST, inferred using the AMR package v3.0.0 [14]. Epidemiological cutoff values (ECOFFs) were used when EUCAST breakpoints were unavailable, with their use indicated in the 'used ECOFF' field as a yes or no entry. Breakpoints for *Salmonella enterica subsp. enterica* were used to determine updated phenotypes for *Salmonella enterica.*

##### S1.7 Data analysis

The origin of CABBAGE data and the overlap between data sources was visualised using ComplexUpset (version 1.3.5) in R. GeoPandas (version 1.1.1) in Python was used to map the global distribution of entries and their resistance phenotypes, restricted to countries with at least 500 isolates linked to a resistance phenotype for the latter. The comparative analysis of CABBAGE with ATLAS was carried out on CABBAGE data that was filtered to entries with a collection year between 2004-2023 to reflect the range of collection dates in ATLAS. An analysis of CABBAGE data that was not filtered by collection year was also performed. The CABBAGE phenotypes derived from the original data source were used for this analysis, rather than the updated phenotypes. Logistic regression in Python was used to assess bias towards resistance in either dataset. Fisher's exact test was applied to identify specific pathogen-antibiotic pairs that showed significant bias towards susceptibility or resistance.

##### S1.8 AMR portal data preparation and execution

Unassembled bacteria genomes were generated using Shovill (v1.1.0) with contigs screened against the T2T CHM13 version 2 assembly (GCA 009914755.4) with MUMmer (v4.0.0rc1) to remove human contamination. As part of the mettannoator annotation (v1.5.0), AMRFinderPlus (v4.0.23) using database version 4.0 2025-07-16.1 was used.

Both phenotype and genotype records were transformed into CSV files using a set of custom python scripts (https://github.com/Ensembl/amr_genotypes), normalised and then converted into a final set of parquet and CSV data files. Antibiotic agents were converted to Antibiotic Resistance Ontology (ARO) terms using EMBL-EBIs Ontology Lookup Service [16] [17]. Fields such as measurement, resistance phenotype and species information were additionally harmonised. The AMR portal (https://github.com/Ensembl/amr-portal/) ETL suite provides a configuration file format to describe data sets, create composite representations and annotate as URLs or similar. After an ETL run a single DuckDB file is generated representing the portal data set and contains all configuration needed. The portal was built using a Python FastAPI backend which creates SQL statements to filter each data set accordingly. The front-end is a single page application using the lit framework.

##### S1.9 Supplementary Results

The inclusion of PubMed IDs and database names in CABBAGE entries allow, when available, users to refer to the original data source. however, this information isn't available for every entry, and where the PubMed ID is recorded, the majority of isolates are associated with a single ID, while others are associated with up to eight (Figure S2).

The number of entries available for each antibiotic is pathogen specific. Consequently, some antibiotics have been tested more frequently against certain pathogens than others, as illustrated in Supplementary Figure S3. For instance, the most frequently tested antibiotics for *M.tuberculosis* are rifampin (12.4%) and isoniazid (12.2%), consistent with their role as first-line treatments. In contrast, amongst the 91 most common antibiotics in CABBAGE (>1,000 entries), eight (erythromycin, fusidic acid, gentamicin, methicillin, mupirocin, penicillin, tetracycline and vancomycin) have only a single AST result each, while 62 have no data for *M.tuberculosis*. Amongst the antibiotics tested against the widest range of pathogens are ciprofloxacin (54/58 pathogens), gentamicin (53/58), meropenem (52/58) and ceftriaxone (52/58), highlighting their use as broad-spectrum antimicrobials. Conversely, 43 antibiotics were only tested against a single pathogen each.

The availability of AST data (either MIC, phenotype, or both) also varies between different antibiotics, as shown in Supplementary Figure S6. For instance, ciprofloxacin is the most frequently tested antimicrobial (>96,000 entries), while dicloxacillin, ceftizoxime and mecillinam have the least data availability, each represented by only a single AST result. Broad-spectrum antibiotics and/or those that are commonly used in clinical settings are, as might be expected, the most frequently represented in CABBAGE, while there is less available data for narrow-spectrum and rarely used antimicrobials.

##### S1.10 Supplementary Tables and Figures

| Search term | Hits | Relevant | Precision |
|---|---|---|---|
| (“antibiotic resistance” OR “antibiotic sensitivity” OR “drug resistance” OR “multidrug resistance” OR AMR OR MDR) AND (”genotype phenotype correlation” OR “genotype phenotype prediction”) AND (”whole genome sequencing” OR WGS) | 84 | 21 | 25% |
| (“antibiotic resistance” OR “antibiotic sensitivity” OR “antimicrobial resistance” OR “antimicrobial sensitivity testing” OR “drug resistance” OR “multidrug resistance” OR AMR OR AST OR MDR) AND (”genotype phenotype correlation” OR “genotype phenotype prediction” OR “genotype phenotype association” OR “genome wide association study” OR GWAS) AND (”next generation sequencing” OR “whole genome sequencing” OR NGS OR WGS) | 197 | 32 | 16% |
| (antibiotic OR antimicrobial OR drug) AND resistance AND sequencing AND accession | 100 | 6 | 6.0% |
| (antibiotic OR antimicrobial OR drug) AND resistance AND sequencing AND prediction | 1205 | 89 | 7.4% |
| predict* NOT prediction AND (antibiotic OR antimicrobial OR drug) AND resistance AND sequencing | 1760 | 43 | 2.4% |
| sequenc* NOT sequencing AND predict* AND (antibiotic OR antimicrobial OR drug) AND resistance | 979 | 16 | 1.6% |

Table S1: Terms used in the literature search, with a summary of their results

| Database | Pathogen | Number of entries |
|---|---|---|
| BV-BRC | *Acinetobacter baumannii* | 21,644 |
| | *Campylobacter coli & jejuni* | 8,301 |
| | *Clostridioides difficile* | 3,990 |
| | *Enterobacter cloacae & hormaechei* | 16,400 |
| | *Enterococcus faecium* | 7,557 |
| | *Escherichia coli* | 172,631 |
| | *Haemophilus influenzae* | 705 |
| | *Helicobacter pylori* | 265 |
| | *Klebsiella pneumoniae* | 109,185 |
| | *Morganella morganii* | 1,399 |
| | *Mycobacterium tuberculosis* | 72,085 |
| | *Neisseria gonorrhoeae* | 45,877 |
| | *Proteus mirabilis* | 1,724 |
| | *Providencia rettgeri & stuartii* | 377 |
| | *Pseudomonas aeruginosa* | 9,839 |
| | *Salmonella enterica* | 151,476 |
| | *Serratia marcescens* | 4,154 |
| | *Shigella flexneri & sonnei* | 18,465 |
| | *Staphylococcus aureus* | 32,441 |
| | *Streptococcus pneumoniae* | 78,087 |
| CDC | *Acinetobacter baumannii* | 1,106 |
| | *Clostridioides difficile* | 180 |
| | *Enterobacter cloacae* | 1,154 |
| | *Enterococcus faecium* | 83 |
| | *Escherichia coli* | 3,027 |
| | *Klebsiella pneumoniae* | 3,716 |
| | *Morganella morganii* | 76 |
| | *Neisseria gonorrhoeae* | 357 |
| | *Proteus mirabilis* | 262 |
| | *Providencia rettgeri & stuartii* | 151 |
| | *Pseudomonas aeruginosa* | 1,919 |
| | *Salmonella enterica* | 71 |
| | *Serratia marcescens* | 316 |
| | *Shigella flexneri & sonnei* | 241 |
| | *Staphylococcus aureus* | 1,895 |
| COMPARE | *Escherichia coli* | 11,214 |
| | *Salmonella enterica* | 21,266 |
| Microreact | *Klebsiella pneumoniae* | 4,319 |
| | *Streptococcus pneumoniae* | 39,418 |
| | *Salmonella enterica* | 1,843 |
| NARMS | *Campylobacter coli & jejuni* | 24,208 |
| | *Salmonella enterica* | 195,248 |
| | *Escherichia coli* | 10,853 |
| | *Shigella sonnei & flexneri* | 9,561 |
| NDARO | *Acinetobacter baumannii* | 10,717 |
| | *Campylobacter coli & jejuni* | 30,487 |
| | *Enterobacter cloacae & hormaechei* | 1,805 |
| | *Enterococcus faecium* | 94 |
| | *Escherichia coli* | 45,750 |
| | *Klebsiella pneumoniae* | 6,714 |
| | *Morganella morganii* | 57 |
| | *Neisseria gonorrhoeae* | 56 |
| | *Proteus mirabilis* | 128 |
| | *Providencia rettgeri & stuartii* | 95 |
| | *Pseudomonas aeruginosa* | 1,681 |
| | *Salmonella enterica* | 114,779 |
| | *Serratia marcescens* | 165 |
| | *Shigella sonnei* | 160 |
| | *Staphylococcus aureus* | 1,533 |
| | *Streptococcus pneumoniae* | 3,506 |
| PathogenWatch | *Neisseria gonorrhoeae* | 56,686 |
| | *Staphylococcus aureus* | 5,278 |
| PubMLST | *Haemophilus influenzae* | 139 |
| | *Staphylococcus aureus* | 24 |
| | *Neisseria gonorrhoeae* | 45,690 |
| | *Neisseria meningitidis* | 2,659 |
| | *Streptococcus pneumoniae* | 464 |

Table S2: The number of entries for each pathogen from each database after processing. Note that these entries may be shared amongst multiple databases.

| Database | Number of entries |
|---|---|
| BV-BRC | 764,007 |
| CDC | 14,657 |
| COMPARE | 32,480 |
| Microreact | 45,580 |
| NARMS | 240,218 |
| NDARO | 217,946 |
| PathogenWatch | 61,964 |
| PubMLST | 48,976 |

Table S3: The number of entries contributed by each database after processing. Entries may be shared amongst multiple databases.

| **Feature** | **Count** |
|---|---|
| Genomes | 170,750 |
| Measurement values | 1,299,368 |
| Phenotypes | 1,379,999 |
| Genera | 20 |
| Antibiotics | 165 |

Table S4: Features of CABBAGE and their corresponding counts. The phenotype count represents the number of phenotypes extracted from the original data source, rather than the updated phenotype (EUCAST or CLSI).

| PubMed ID | Laboratory Typing Method |
|---|---|
| 22914622 | broth dilution |
| 23299977 | broth dilution |
| 24277043 | E-test |
| 24462211 | agar dilution |
| 25378573 | agar dilution |
| 25903077 | disk diffusion |
| 26135860 | broth dilution |
| 26806258 | disk diffusion |
| 26935729 | agar dilution |
| 26974227 | E-test |
| 27067331 | broth dilution |
| 27150362 | disk diffusion |
| 27322919 | broth dilution |
| 27432602 | E-test |
| 28931025 | disk diffusion |
| 29017452 | broth dilution |
| 29295910 | disk diffusion |
| 29377579 | agar dilution |
| 29547094 | broth dilution |
| 29553335 | agar dilution (SPT) and E-test (all other antibiotics) |
| 29617860 | broth dilution |
| 29701830 | disk diffusion (PEN,TCY) and E-test (all other antibiotics) |
| 29723570 | E-test |
| 29882175 | E-test |
| 30820563 | agar dilution |
| 31005733 | broth dilution |
| 31010866 | broth dilution |
| 31038449 | broth dilution |
| 31220112 | E-test |
| 31253105 | disk diffusion (AMP, CHL, SXT) and broth dilution (all other antibiotics) |
| 31358980 | E-test |
| 31424550 | E-test |
| 31488838 | agar dilution |
| 31504566 | E-test |
| 31530672 | broth dilution |
| 31665255 | E-test |
| 31941492 | broth dilution (COL, DOX,PLB) and disk diffusion (all other antibiotics) |
| 31978353 | agar dilution |
| 32010104 | disk diffusion |
| 32013864 | E-test |
| 32048461 | agar dilution |
| 32068837 | E-test |
| 32091356 | agar dilution |
| 32205351 | broth dilution |
| 32422315 | disk diffusion |
| 32829411 | E-test |
| 32868325 | E-test |
| 32977759 | E-test |
| 33367806 | E-test |
| 34161790 | agar dilution |
| 34570642 | broth dilution |
| 34693903 | broth dilution |
| 34745054 | broth dilution |
| 34819373 | broth dilution |
| 35156028 | broth dilution |
| 35178041 | broth dilution |
| 35260883 | broth dilution |
| 35308374 | agar dilution |
| 35378228 | broth dilution |
| 35651495 | broth dilution |
| 35731173 | broth dilution |
| 35880764 | agar dilution |
| 35916524 | broth dilution |
| 36165686 | broth dilution |
| 36208202 | agar dilution |
| 36473954 | broth dilution |
| 36801013 | broth dilution |
| 37316492 | broth dilution |
| 37327220 | E-test (CIP, AZM) and disk diffusion (all other antibiotics) |
| 37549252 | disk diffusion |
| 3805277 | broth dilution |

Table S5: PubMed ID's of datasets obtained from BV-BRC and the corresponding laboratory typing methods, determined by referring to the original manuscript and Supplementary Materials.

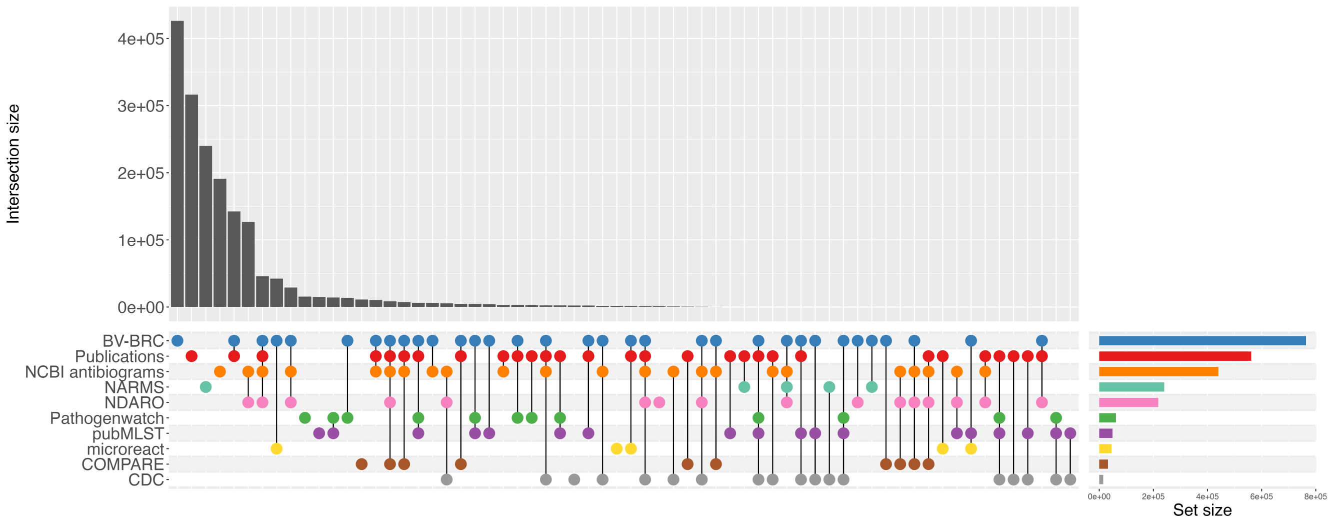


Figure S1: An UpSet plot illustrating the origin of data within the CABBAGE dataset, and the overlap between those sources. Data was sourced from publications, BV-BRC, NCBI antibiograms, NARMS, NDARO, Pathogenwatch, PubMLST, Microreact, COMPARE and the CDC.

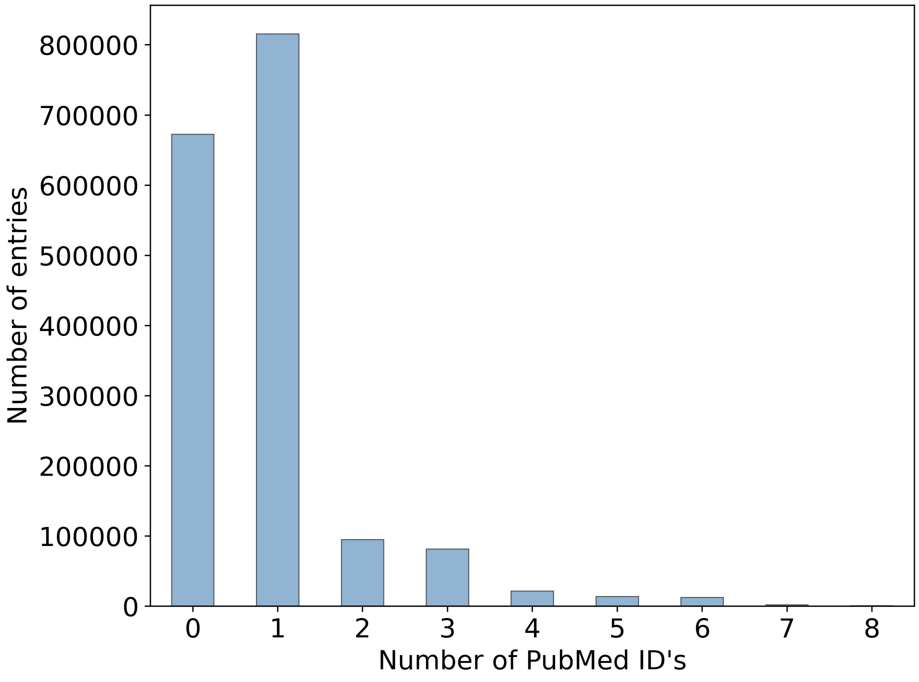


Figure S2: The number of entries associated with multiple, a single, or no PubMed ID.

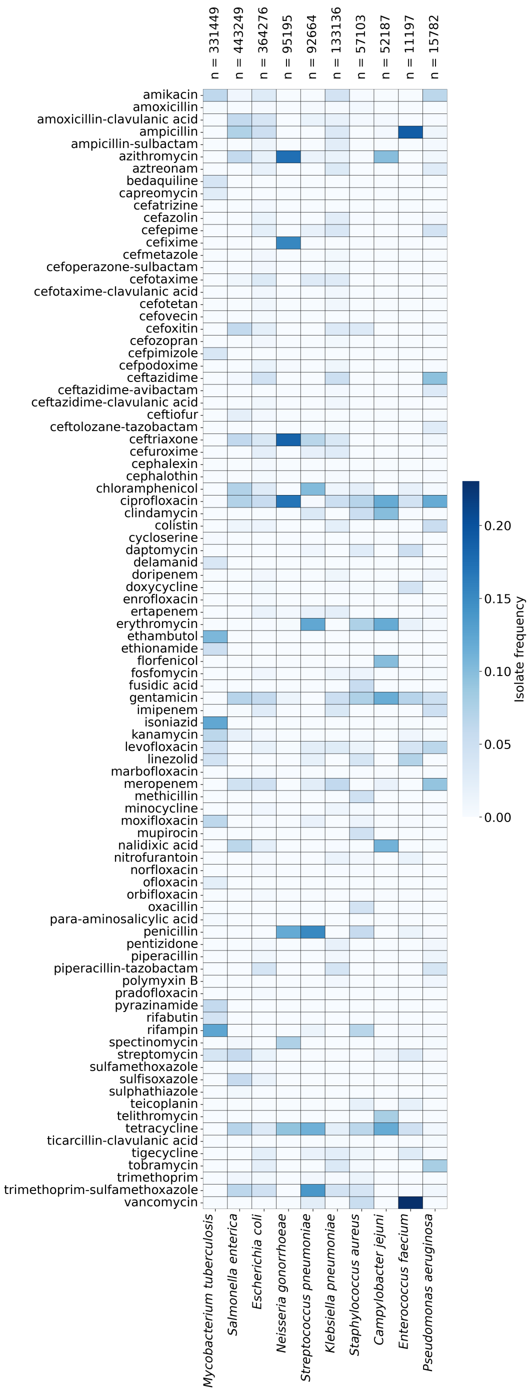


Figure S3: Heat map showing the antibiotic testing frequency (either measurment value, phenotype, or both) across the 10 most common bacterial species in CABBAGE (by unique BioSample ID rather than number of entries). The colour of each cell represents the testing frequency for a specific pathogen-antibiotic pair, with dark blue indicating a high testing frequency and light blue indicating antibiotics that were rarely tested against the corresponding pathogen. The data was restricted to antibiotics represented by >=1,000 entries in CABBAGE.

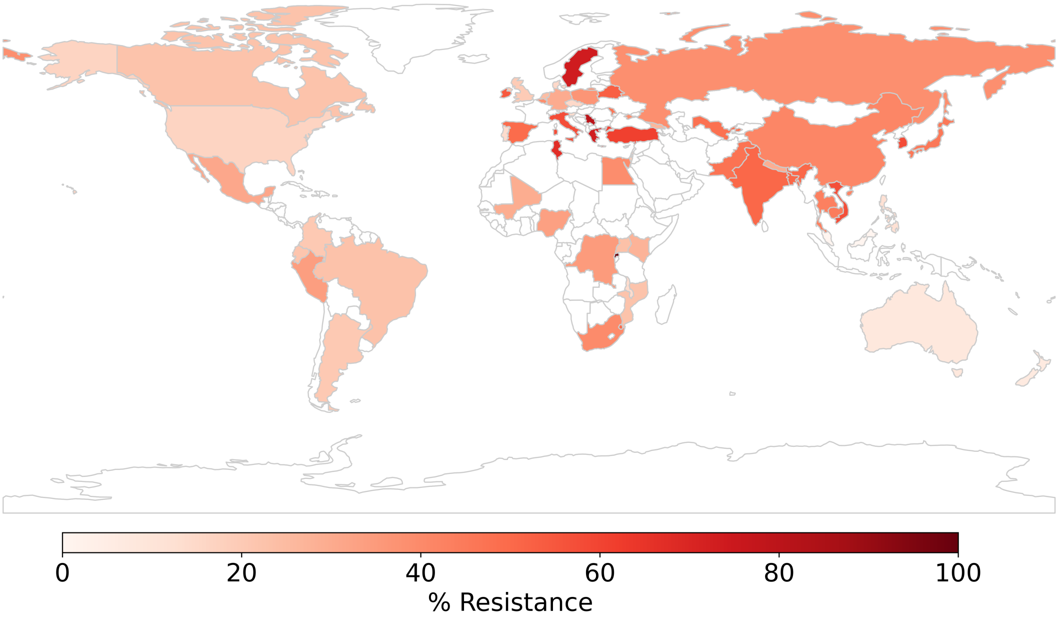


Figure S4: Antimicrobial resistance frequencies by country among phenotypically characterised isolates. Phenotypes derived from the original data source are shown, and countries with fewer than 500 entries are shown in white.

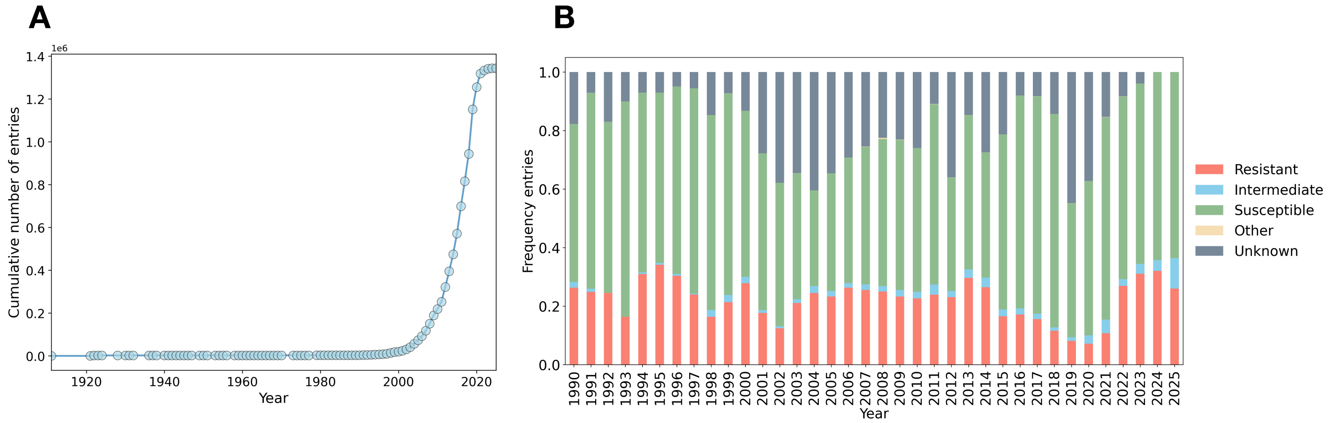


Figure S5: A) The cumulative number of entries (isolate-antibiotic pairs) over time (years), where the isolate collection date was available. B) The frequency of isolate-antibiotic pairs associated with each phenotype between 1990-2025. The category 'other' includes susceptible-dose dependent and non-susceptible.

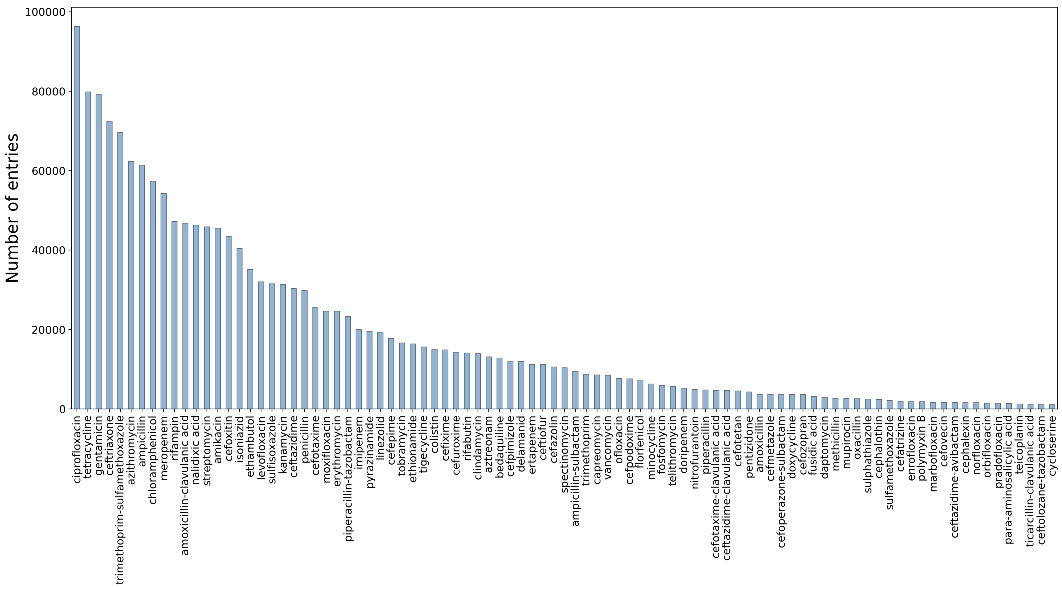


Figure S6: The number of isolates tested against each antibiotic. The data was restricted to antibiotics represented by $\geq$ 1,000 isolates in CABBAGE.

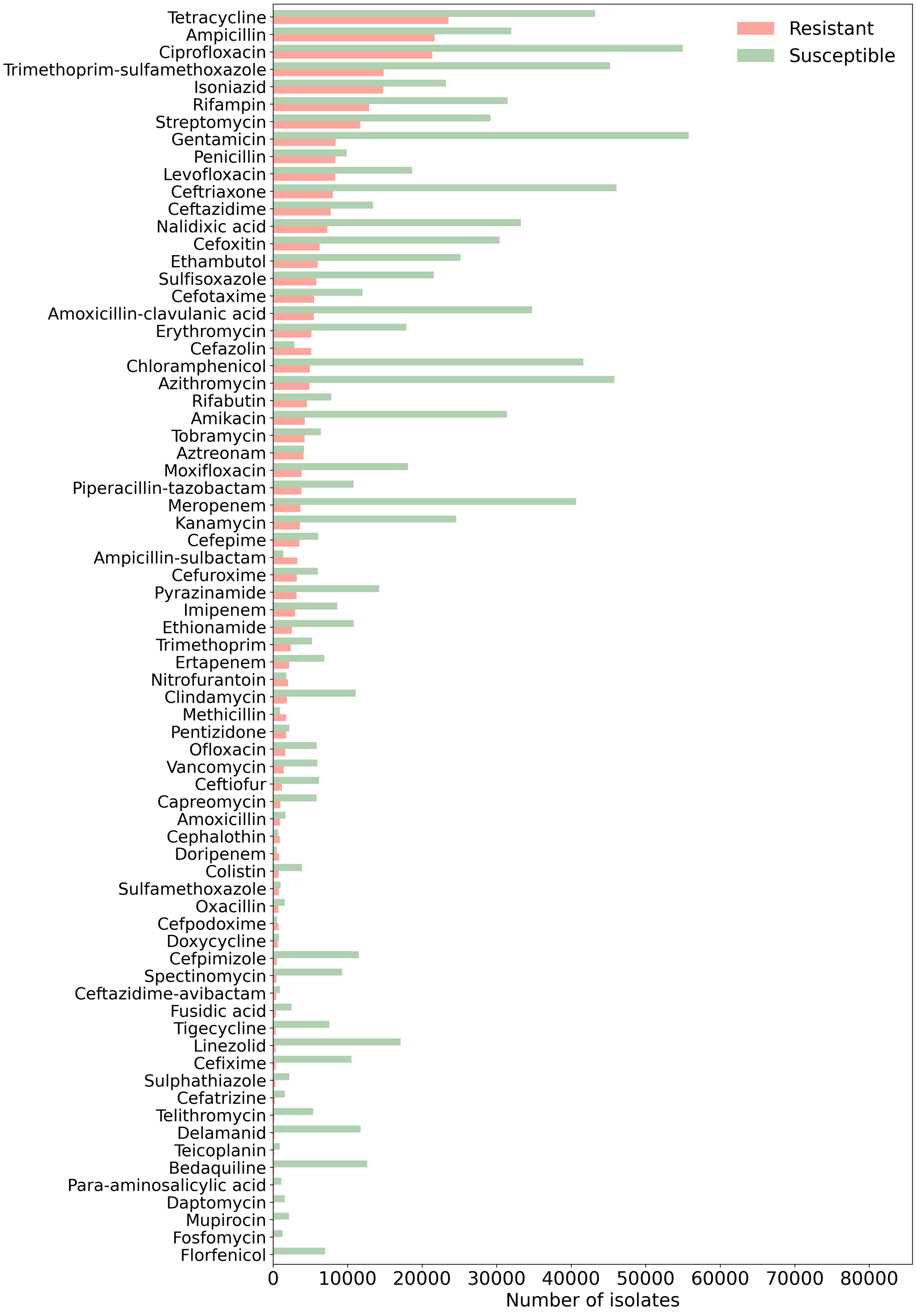


Figure S7: The number of resistant (red bars) versus susceptible (green bars) phenotypes is shown for each antibiotic. The data was restricted to those antibiotics with $\geq 1{,}000$ isolates in CABBAGE.

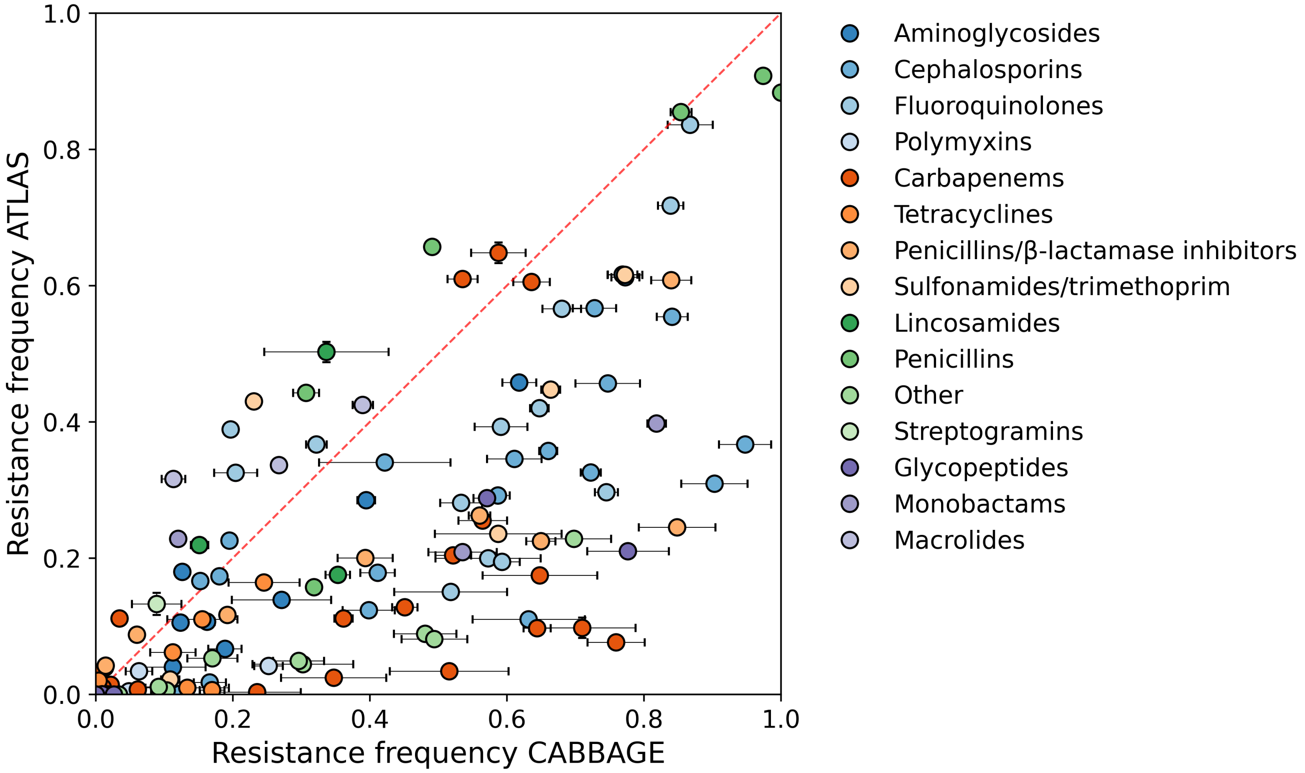


Figure S8: A scatter plot showing the frequency of resistance in CABBAGE and ATLAS, with each data point representative of a specific pathogen-antibiotic pair that is present in both datasets. The data has not been filtered by year, and only pathogen-antibiotic pairs with at least 100 entries in both CABBAGE and ATLAS are shown. Error bars represent 95% CI and colours are representative of different antibiotic classes. Note that the CABBAGE phenotypes presented here represent the phenotypes derived from the original datasets, rather than updated phenotypes.
